# Activation and self-inactivation mechanisms of the cyclic oligoadenylate-dependent CRISPR ribonuclease Csm6

**DOI:** 10.1101/2020.02.19.952960

**Authors:** Carmela Garcia-Doval, Frank Schwede, Christian Berk, Jakob T. Rostøl, Ole Niewoehner, Oliver Tejero, Jonathan Hall, Luciano A. Marraffini, Martin Jinek

**Affiliations:** Department of Biochemistry, University of Zurich, Winterthurerstrasse 190, CH-8057 Zurich, Switzerland; Biolog Life Science Institute GmbH & Co. KG, Flughafendamm 9a, D-28199 Bremen, Germany; Department of Chemistry and Applied Biosciences, Institute for Pharmaceutical Sciences, ETH Zurich, Vladimir-Prelog-Weg 1-5/10, CH-8093 Zurich, Switzerland; Laboratory of Bacteriology, The Rockefeller University, 1230 York Avenue, New York, New York 10065-6399, USA; Howard Hughes Medical Institute, The Rockefeller University, 1230 York Avenue, New York, NY 10065, US

**Author notes:** Correspondence should be addressed to M.J.

## Abstract

Upon target RNA recognition, type III CRISPR-Cas systems produce cyclic oligoadenylate second messengers to activate downstream effectors including Csm6-family ribonucleases via their CARF domains. Here we show that *Enteroccocus italicus* Csm6 (EiCsm6) degrades its cognate cyclic hexa-AMP (cA6) activator and report the crystal structure of EiCsm6 bound to a cA6 mimic. The structure, combined with biochemical and in vivo functional assays, reveal how cA6 recognition by the CARF domain activates the Csm6 HEPN domains for collateral RNA degradation, and how CARF domain-mediated cA6 cleavage provides an intrinsic off-switch to limit Csm6 activity in the absence of ring nucleases. These mechanisms facilitate rapid invader clearance and ensure termination of CRISPR interference to limit self-toxicity.

## Introduction

CRISPR-Cas systems are defense mechanisms present in many bacteria and most archaea that provide RNA-guided immunity against genetic invaders such as bacteriophages and plasmids^1,2^. CRISPR-Cas systems can be divided in two classes and six types based on their composition and the presence of specific CRISPR-associated (*cas*) genes. Type III systems are characterized by the presence of the signature protein Cas10 in their multisubunit CRISPR RNA (crRNA)-guided effector complexes, termed Csm complexes in subtypes III-A and III-D, and Cmr complexes in subtypes III-B and III-C^3^. These complexes directly recognize invader-derived RNA transcripts and harbor two target-dependent nuclease activities, whereby Cas10 subunit (also known as Csm1 or Cmr2) is allosterically activated to degrade single-stranded DNA in a sequence non-specific manner using its HD domain, while the Csm3/Cmr4 subunits cleave the bound RNA in a sequence specific manner^4-10^.

Additionally, target RNA binding activates the Palm domain of the Cas10 subunit of the Csm/Cmr complexes to convert ATP into cyclic oligoadenylate (cOA) second messengers of various sizes, which in turn allosterically activate CRISPR-associated Rossman-fold (CARF) domain-containing effector proteins including Csm6/Csx1 ribonucleases^11-14^. The type III CRISPR-Cas systems found in different organisms have been shown to produce different cOA species; whereas *Streptococcus thermophilus* and *Enterococcus italicus* generate cyclic hexa-AMP (cA6), *Thermus thermophilus, Sulfolobus islandicus* or *Sulfolobus solfataricus* produce cyclic tetra-AMP (cA4) ^11-14^. Upon activation, the RNase activity of Csm6/Csx1 enzymes, catalyzed by their ‘higher eukaryotes and prokaryotes nucleotide binding’ (HEPN) domains, provides an auxiliary interference mechanism to facilitate invader clearance and may induce dormancy or cell death. Previous studies demonstrated its requirement for efficient in vivo immunity, specifically when targeting late genes^13,15-17^. In some type III CRISPR-Cas systems, cOA signaling is terminated by standalone CARF domain-containing ring nucleases, proteins able to degrade cA4 to yield linear adenosine tetra-and dinucleotides^18^. Furthermore, recent studies have shown that some Csm6-family enzymes can also intrinsically catalyze cOA degradation and thus possess a self-inactivation mechanism^19,20^.

Although Csm6/Csx1 proteins that are activated by cA4 have been recently characterized^20,21^, the molecular mechanisms of cA6-dependent Csm6 RNases remain elusive. In this study, we show that the type III CRISPR-associated RNase Csm6 from *Enterococcus italicus* (EiCsm6), which is activated by cA6, possesses an intrinsic cA6 degradation activity mediated by its CARF domain. Our crystal structure of EiCsm6 in complex with a non-degradable cA6 mimic, together with supporting biochemical and functional studies, reveal the mechanisms of cA6 sensing and degradation, as well as cA6-mediated allosteric activation of EiCsm6. These results provide a mechanistic framework for understanding the function of CARF domain-containing effector RNase in the context of CRISPR-Cas immunity.

## Results

### The CARF domain of Csm6 is a ring nuclease

We previously showed that the ribonuclease Csm6 associated with the type III CRISPR-Cas system from *Enterococcus italicus* is activated by cyclic hexa-AMP (cyclic hexa-adenylate, cA6) ^22^. To test whether EiCsm6 is capable of degrading cA6, we analyzed reaction mixtures containing cA6 and wild-type (WT) EiCsm6 using liquid-chromatography-mass spectrometry (LC-MS), revealing that cA6 was efficiently degraded to yield a mixture of products, including a species with a m/z ratio of 986.1, matching that of linear tri-AMP carrying a 2’,3’-cyclic phosphate group (**Supplementary Figure 1**). cA6 was also efficiently degraded in the presence of RNase-deficient Csm6 containing a point mutation in the HEPN active site (H377A), excluding the possibility that the degradation is catalyzed solely by the HEPN domain (**Supplementary Figure 1**). Corroborating these results, a cA6 sample pre-incubated in the presence of WT EiCsm6 protein was almost inactive in stimulating the RNase activity of WT EiCsm6 in a fluorogenic ribonuclease activity assay, indicative of cA6 degradation (**Supplementary Figure 2a,b**). Preincubation of cA6 with EiCsm6 containing inactivating mutations in the HEPN domain (R372A/N373A/H377A, herein referred to as dHEPN) resulted in partial degradation of cA6 and reduced activation of WT EiCsm6, as did pre-incubation with with EiCsm6 containing a mutation (T114A) in the conserved CARF domain motif. In contrast, preincubation with EiCsm6 containing mutations both in the HEPN RNase active site and in the CARF domain did not result in cA6 degradation (**Supplementary Figure 2b**). Together, these experiments indicate that although the RNase activity of the HEPN domain contributes to cA6 degradation, the CARF domain of EiCsm6 is intrinsically capable of catalyzing cA6 degradation.

### Crystal structure of Csm6 bound to cA6 mimic

The catalytic mechanism of cOA degradation by ring nucleases proceeds through a nucleophilic attack of the ribose 2’-hydroxyl to generate a 2’,3’-cyclic phosphate product^19^. Consistent with this, 2’,2’’,2’’’,2’’’’,2’’’’’,2’’’’’’-hexa-F-c-hexa-dAMP (cFA6), a cA6 mimic containing 2’-fluoro modifications in all positions, was resistant to degradation yet still capable of allosterically activating EiCsm6, albeit more weakly than cA6 (**Supplementary Figure 3a– c**). The half-maximal concentration (EC_50_) of cFA6 was determined to be 170 nM, approximately 50-fold higher than that of cA6^11^, likely due to weaker binding resulting from the 2’-fluoro modifications (**Supplementary Figure 3d**).

To gain structural insights into the mechanism of cA6 binding, we co-crystallized EiCsm6 with cFA6 and determined the structure of the complex at a resolution of 2.4 Å (**Supplementary Table 1**). The structure reveals that EiCsm6 adopts the canonical homodimeric architecture of Csm6/Csx1 enzymes, with extensive dimerization interfaces between the CARF and HEPN domains. A single molecule of cFA6 is bound within a conserved pocket spanning the dimer interface of the CARF domains (**Fig. 1, Supplementary Figure 4a**), as predicted by earlier structural studies of *Thermus thermophilus* Csm6^23^, and recent structures of *Thermococcus onnurineus* (ToCsm6) and *Sulfolobus islandicus* (SisCsx1) proteins bound to c-tetra-AMP^20,21^. The cFA6 ligand exhibits 2-fold rotational symmetry coincident with the 2-fold symmetry axis of the EiCsm6 homodimer, with nucleotides A2-A4 binding to one EiCsm6 protomer, and A1 and A5-A6 to the other, and is surrounded by CARF domain loops comprising residues 10-21, 39-41, 76-82, 111-116, and 137-139 (**Supplementary Figure 4b**). The 2’-fluoro ribose moieties adopt the C2’-endo conformation in all positions, while the adenine bases adopt the *syn* conformation in nucleotides A1/A4 and A2/A5, and *anti* in A3/A6. Structural comparisons of the EiCsm6-cA6, ToCsm6-cA4 and SisCsx1-cA4 complexes indicate that nucleotides A1/A4 and A3/A6 in the EiCsm6 structure are accommodated similarly to their respective counterparts in the ToCsm6 and SisCsx1 structures, binding in structurally analogous (but only partially conserved) pockets in the CARF domains, while nucleotides A2/A5 occupy distinct sites at the CARF domain interface (**Supplementary Figure 5**).

**Figure 1.**
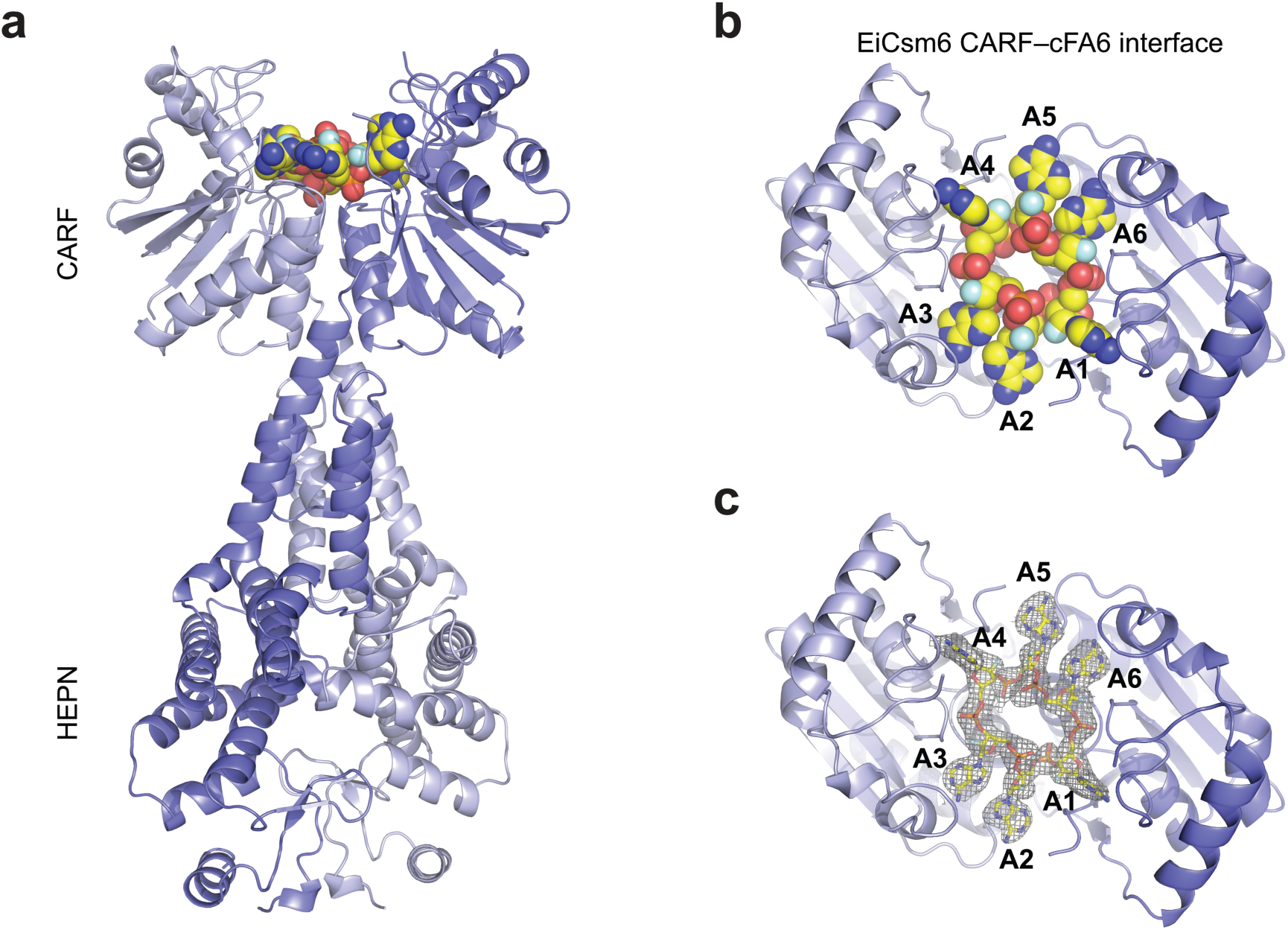
Cyclic hexaadenylate recognition by EiCsm6. (**a, b**) Overall structure of EiCsm6 homodimer bound to cFA6, shown in two orientations (a: side view of the EiCsm6 homodimer; b: view of the cFA6 binding site at the CARF domain interface). The EiCsm6 homodimer is shown in cartoon representation (with protomers colored light and dark blue). cFA6 is depicted in sphere format. (**c**) Zoom-in view of the cA6 binding site at the CARF domain interface of EiCsm6, showing a 2m*F*_O_-D*F*_C_ composite omit map, contoured at 1.0 σ and displayed within a radius of 2.2 Å around cFA6.

### cA6 sensing by the Csm6 CARF domain

The EiCsm6-cFA6 structure reveals that the ligand is recognized by an extensive network of ionic and hydrogen bonding interactions with both the ribose-phosphate backbone and the adenine bases (**Supplementary Figure 4b, Supplementary Figure 6**). The bases of nucleotides A1/A4 are inserted into a deep pocket and are recognized via hydrogen-bonding interactions with the side chain of Arg173 and the backbone carbonyl of Asp19, while A2/A5 and A3/A6 interact with the side chains of Glu80 and Thr39/Ser41, respectively (**Fig. 2a,b**). In turn, Gln116 forms a hydrogen bond with the phosphodiester group connecting A2/A5 and A3/A6 (**Fig. 2c**). Alanine substitution of a subset of conserved residues involved in cA6 binding, including Asn10, Thr114 and Gln116 (**Supplementary Figure 6**), resulted in the reduction or near-complete loss of cA6-dependent RNase activity, confirming the importance of these residues for cA6 sensing (**Fig. 2d**).

**Figure 2.**
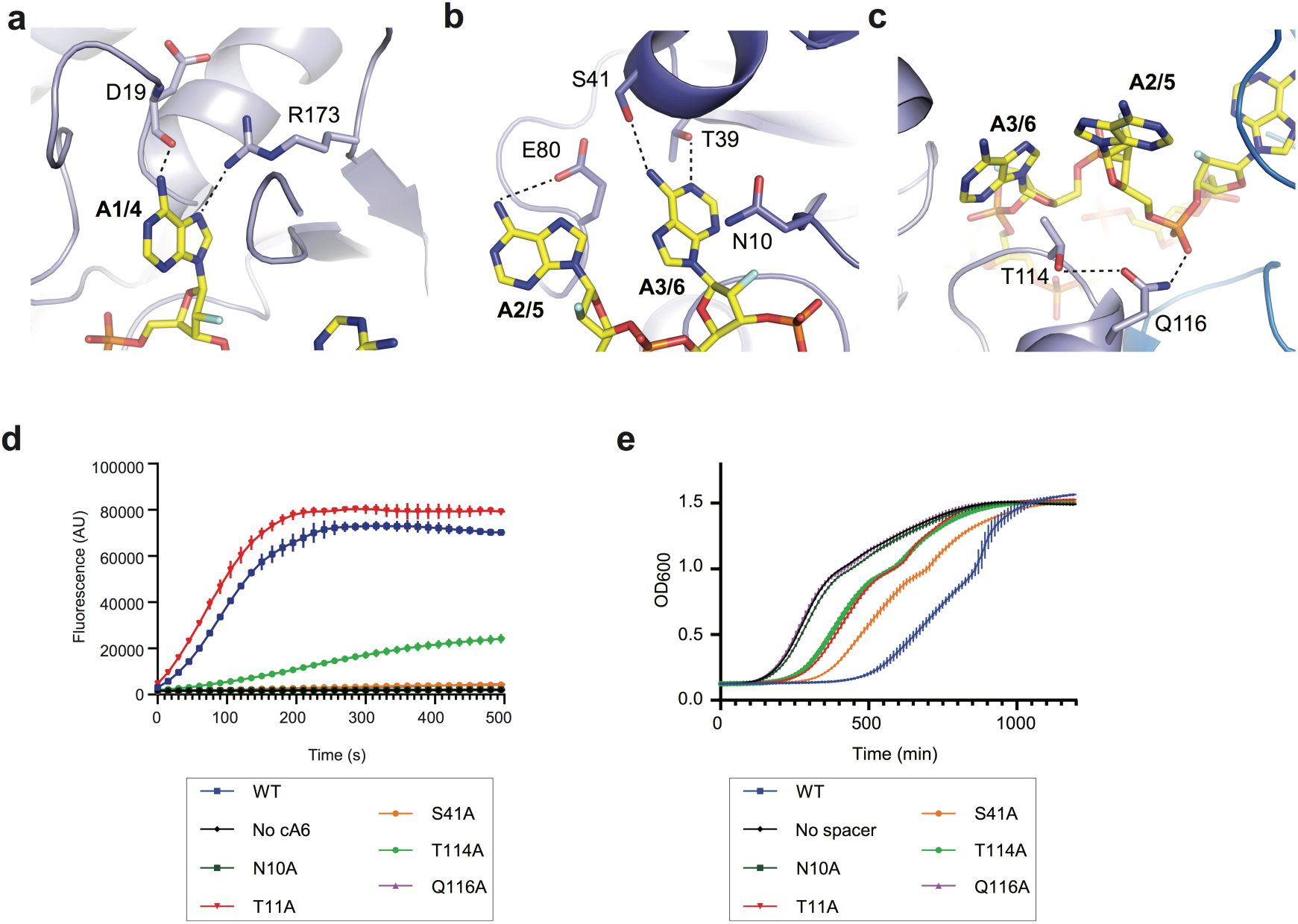
The CARF domain of EiCsm6 recognizes cA6. (**a, b, c**) Protein-cFA6 interactions. EiCsm6 is depicted in cartoon representation; cFA6 is shown as sticks. Hydrogen bonding interactions are represented as black dotted lines. (**d**) Fluorogenic RNase activity assay of CARF domain mutants of EiCsm6 (0.5 nM) in the presence of 10 nM cyclic-hexa-AMP (cA6). AU, arbitrary units. (**e**) Growth curves (optical density at a wavelength of 600 nm) of *Staphylococcus aureus* strains harboring pTarget and pCRISPR plasmids. The pCRISPR plasmid expresses wild-type or CARF domain mutants of EiCsm6 and the *Staphylococcus epidermidis* Cas10-Csm effector complex containing an inactivating point mutation in the HD domain of Cas10. Data points in panels d and e represent the mean of three replicates; error bars represent the standard error of the mean (s.e.m.). Source data are provided as a Source Data file.

In previous studies, we showed that the RNase activity of Csm6 in vivo depends on its allosteric activation by cyclic oligoadenylates and demonstrated the compatibility between EiCsm6 and the *Staphylococcus epidermidis* Type III Cas10-Csm complex^22^. To validate our structural insights into cA6 sensing, we tested the in vivo activity of wild-type and CARF domain mutant EiCsm6 proteins when co-expressed with the *S. epidermidis* type III CRISPR-Cas system lacking DNase activity in the HD domain of the Cas10 subunit (dHD) and an inducible RNA target. Target-induced activation of WT EiCsm6 triggered growth arrest due to its collateral RNase activity, which was dependent on the presence of a targeting spacer in the CRISPR array, as shown previously for *S. epidermidis* Csm6^17^. In contrast, EiCsm6 mutants in which specific residues involved in cA6 sensing were substituted with alanine either did not cause growth arrest or impeded growth only weakly, thus validating the importance of the CARF domain residues for cA6 sensing and Csm6 activation in vivo (**Fig. 2e**).

### Mechanism of autocatalytic cA6 degradation

The EiCsm6-cFA6 structure provides insights into the mechanism underpinning cA6 degradation. As a consequence of the C2’-endo ribose pucker in A6/A3 and the positioning of A1/A4 in its binding pocket, the A6-A1 (and A3-A4) dinucleotide is forced into a conformation favoring an in-line nucleophilic attack of the A6/A3 2’-hydroxyl group onto the downstream scissile phosphodiester group to form a 2’,3’-cyclic phosphate trinucleotide product (**Fig. 3a**). Whereas the 2’-fluoro group of A6/A3 (and, presumably, the 2’-hydroxyl in cA6) is in hydrogen-bonding contact with the backbone amide group of Asn10, the scissile phosphodiester group is stabilized by hydrogen-bonding interactions with the backbone amide group and the side chain of Thr11. Thus, cA6 degradation by EiCsm6 seems to be mediated primarily by steric factors that force the ligand to adopt a conformation compatible with in-line nucleophilic substitution and does not involve deprotonation of the attacking nucleophile or protonation of the leaving group by acid-base catalysis, as postulated for ring nucleases and other Csm6/Csx1 enzymes in previous studies^18-20^. To verify the functional role of Thr11 in cA6 degradation, we compared the cA6-dependent RNase activities of WT and T11A EiCsm6 proteins. The T11A EiCsm6 protein retained the ability to be allosterically activated by cA6 in vitro (**Fig. 2d**), although the mutation caused only a weak growth arrest phenotype in vivo (**Fig. 2e**), presumably because the cA6 levels generated by the Cas10-Csm complex upon target induction were insufficient to activate T11A EiCsm6 due to partially compromised cA6 binding. The T11A EiCsm6 protein was impaired in cA6 degradation, as indicated by a pre-incubation experiment (**Supplementary Figure 7a**) and confirmed by LC-MS analysis (**Supplementary Figure 7b**). Additionally, the T11A EiCsm6 protein displayed sustained in vitro RNase activity in the presence of 500 nM cA6, as compared to WT Csm6, further indicating that it is deficient in cA6 degradation (**Fig. 3b**). Collectively, these results suggest that cA6 degradation by the CARF domain of EiCsm6 intrinsically limits the sequence non-specific RNase activity of the enzyme’s HEPN domain, which might prevent self-toxicity and contribute to the termination of the CRISPR immune response in vivo. To test this hypothesis, we examined the effect of expressing WT or T11A EiCsm6 in the presence of a *S. epidermidis* Cas10-Csm complex that was deficient in target RNA degradation due to an inactivating mutation in the Csm3 subunit (D32A, dCsm3), which results in sustained production of cA6. At low levels of target induction, WT EiCsm6 was unable to cause growth arrest (**Fig. 3c**). In contrast, the T11A EiCsm6 mutation resulted in a strong growth inhibition phenotype, indicating that the loss of ring nuclease activity in the CARF domain results in the hyperactivation of EiCsm6.

**Figure 3.**
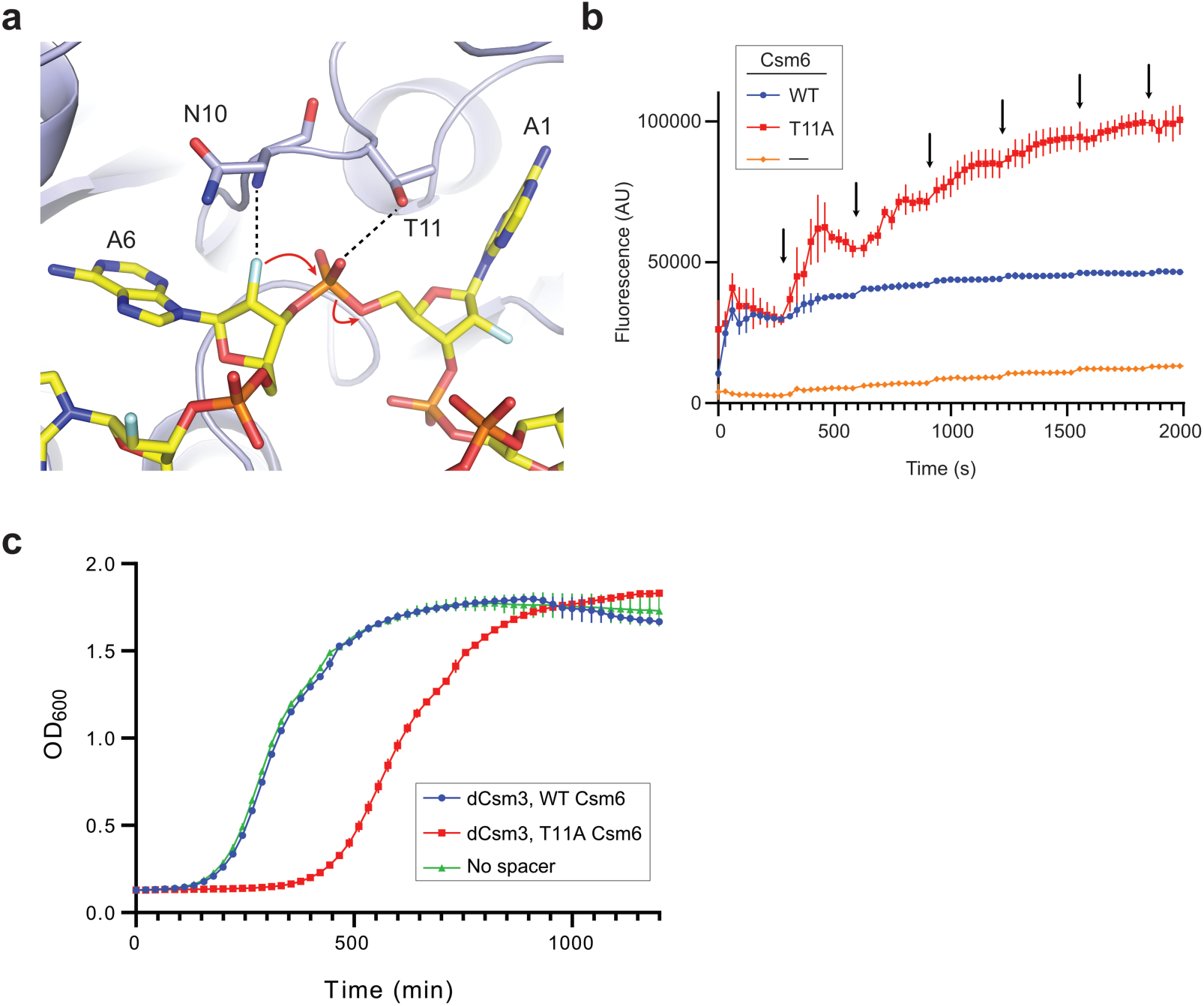
The CARF domain of EiCsm6 catalyzes cA6 hydrolysis. (**a**) Zoom-in view of nucleotides A1 and A6 of cFA6 bound to the EiCsm6 CARF domain. Residues implicated in the cleavage are represented as sticks; hydrogen bonding interactions are depicted as black dotted lines. The red arrows represent the proposed reaction mechanism of cA6 cleavage involving nucleophilic attack of the 2’-hydroxyl group of A6 onto the phosphodiester group connecting A6 and A1. (**b**) RNase activity assay comparing the activity of WT and T11A EiCsm6 proteins in the presence of 500 nM cA6. The black arrows represent spike-ins of 0.8 pmol of RNaseAlert substrate at 5 min intervals. AU, arbitrary units. (**c**) Growth curves (optical density at a wavelength of 600 nm) of *Staphylococcus aureus* strains harboring pTarget and pCRISPR plasmids. The pCRISPR plasmid expresses wild-type or CARF domain mutants of EiCsm6 and the *Staphylococcus epidermidis* Cas10-Csm effector complex containing an inactivating point mutation in the Csm3 subunit (D32A; dCsm3). Data points in panels b and c represent the mean of three replicates; error bars represent the standard error of the mean (s.e.m.). Source data are provided as a Source Data file.

### cA6-dependent activation mechanism of Csm6

Owing to the homodimeric architecture of Csm6 enzymes, their HEPN nuclease domains form a composite RNase active site at the domain interface^23^. In the EiCsm6-cFA6 structure, the site is occupied by a sulfate ion originating from the crystallization solution, acting as a mimic of the scissile phosphate in an RNA substrate (**Fig. 4a**). The anion is contacted by salt-bridge interactions with Arg372 and flanked by a pair of His377 side chains, which are part of the R-X_4_-H HEPN domain motif and predicted to elicit acid-base catalysis based on homology with other HEPN-domain RNases^24^ (**Supplementary Figure 8**). The disposition of the active site residues is similar as in the activated-state structures of both RNase L and cA4-recognizing Csm6/Csx1 RNases ToCsm6 and SisCsx1^20,21,25,26^, indicating that the HEPN domains adopt the activated conformation in the EiCsm6-cFA6 complex.

**Figure 4.**
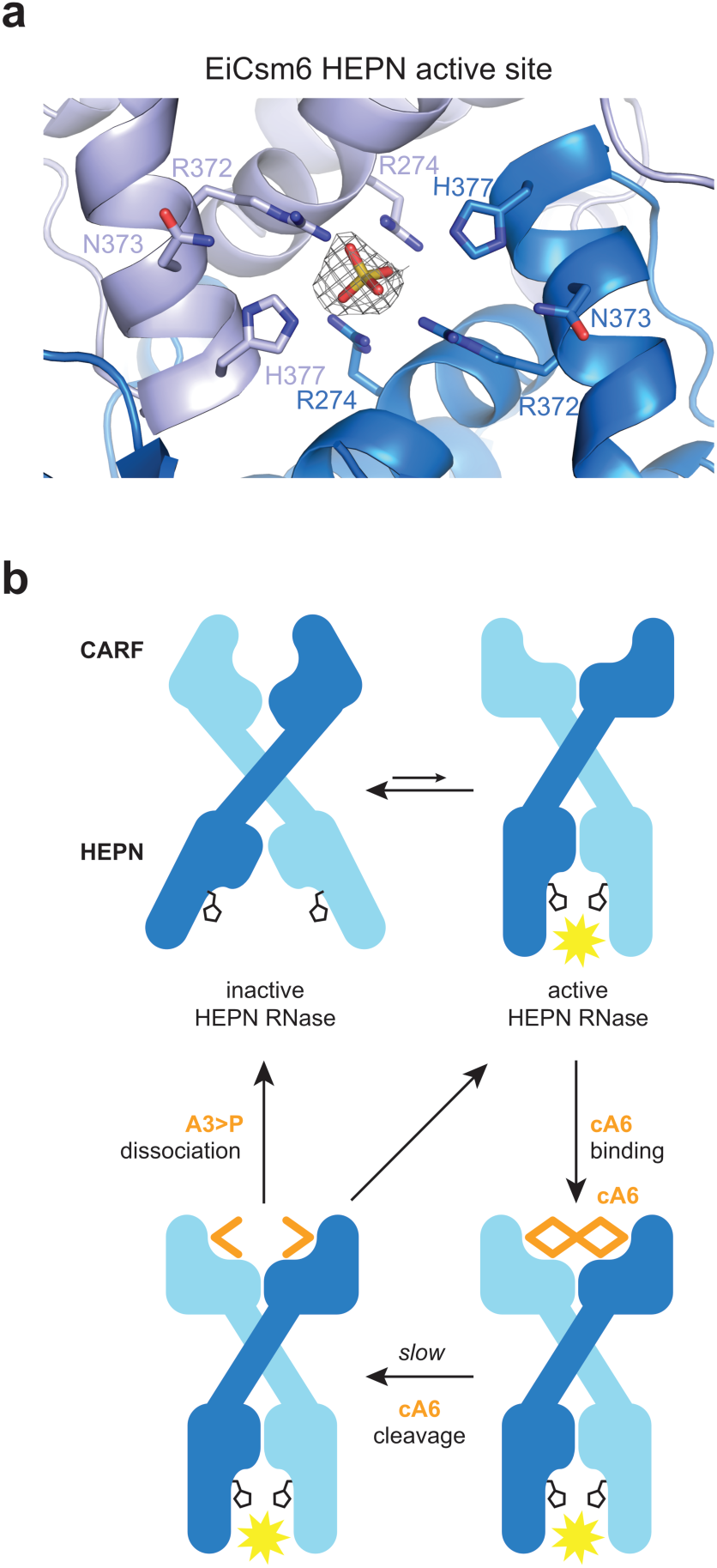
Model of cA6-mediated allosteric activation mechanism of Csm6 Rnases. (**a**) Zoom-in view of the HEPN domain ribonuclease active site in EiCsm6. The conserved R372, N373, H377 residues of the HEPN catalytic motif, and additionally R274, are represented as sticks. Electron density, interpreted as a sulfate ion (depicted as 2m*F*_O_-D*F*_C_ composite omit map, contoured at 1.0 σ and displayed within a radius of 2.2 Å), is located within hydrogen bonding distance of H377, likely mimicking the scissile phosphate group of an RNA substrate. (**b**) Schematic model of the cA6-mediated allosteric activation mechanism of Csm6 enzymes.

Some Csm6 enzymes display basal HEPN-dependent RNase activity in the absence of cOA ligands^15,23,27^, suggesting that their HEPN domains sample a conformational equilibrium between activated and inactive states. This is supported by the observation that the conformations of apo-ToCsm6 and apo-SisCsx1 are nearly identical to their respective cA4-bound structures^20,21^, suggesting that they may represent the activated state. Based on these considerations, an allosteric activation model for EiCsm6 can be envisioned (**Fig. 4b**). In the absence of cA6, EiCsm6 exists in a dynamic conformational equilibrium between inactive and activated forms, with the inactive form(s) being predominant. The HEPN domain active center and the CARF domain cA6 binding site are allosterically coupled. cA6 binding to the CARF domain interface preferentially stabilizes the activated form, in which the HEPN domain histidine residues are properly oriented to catalyze RNA degradation by an acid-base mechanism. While bound to cA6, the CARF domains act as a ring nucleases whose slow activity converts cA6 into two molecules of 2’,3’-cyclic phosphate-terminated tri-AMP and triggers product dissociation. This leads to termination of the HEPN domain RNase activity by destabilization of the activated conformation. The biological activity of Csm6 enzymes is thus controlled by the interplay of the HEPN domain RNase activity and the slow, *cis*-acting ring nuclease activity of the CARF domain.

## Discussion

In sum, the structural and biochemical studies presented in this study provide insights into the molecular mechanisms responsible for cA6 sensing, allosteric activation, and autocatalytic inactivation in Csm6 enzymes. We previously demonstrated the allosteric activation of EiCsm6 by cA6 generated by the Palm domain of Cas10 upon RNA target recognition^22^. We now show that the CARF domain of EiCsm6 responsible of cOA sensing also catalyzes the degradation of the substrate, acting as an intrinsic timer that limits the non-specific RNase activity of the HEPN domain. The generation of cA6 degradation products in the presence of a HEPN-domain catalytic mutation proves that the CARF domain of EiCsm6 harbors a ring nuclease activity, although the RNase activity of the HEPN domain also contributes to cOA degradation, as suggested by other studies^20^. By degrading cOA second messengers in *cis*, the ring nuclease activity of Csm6 enzymes thus complements the RNase activity of the Cas10 effector complex that mediates target RNA degradation, which results in the termination of cOA synthesis by the Cas10 Palm domain.

The structure of EiCsm6 bound to cFA6 reveals the cOA recognition and degradation mechanisms. The homodimeric architecture of EiCsm6 results in the formation of a high-affinity cA6 binding site at the CARF domain interface. Based on the structure, we identify several residues involved in cA6 recognition and confirm their role in second messenger sensing in vitro and in vivo. We additionally show that perturbation of the ring nuclease activity of the CARF domain (by targeting the CARF domain residue Thr11 involved in transition state stabilization) results in a hyperactive Csm6 RNase with a slower inactivation profile. The high degree of conservation of the residues involved in cOA binding and degradation suggests that our proposed mechanisms likely apply to other cA6-specific Csm6 proteins (**Supplementary Figure 6**). Notably, the cA6 degradation mechanism of EiCsm6 does not involve acid-base catalysis, contrary to mechanisms postulated for other Csm6/Csx1 enzymes and ring nucleases^18-20^. Finally, the superposition of the EiCsm6 HEPN domains to other HEPN-domain nucleases indicates that EiCsm6 bound to cFA6 is stabilized in an activated state, pointing to a mechanism for cOA-dependent allosteric activation of Csm6/Csx1 proteins relying on conformational selection.

In the context of CRISPR-Cas immunity, self-degradation of cOA by Csm6/Csx1-family RNases likely provides a mechanism for temporal control of their collateral RNase activity. Together with the target-directed RNase activity of the Cas10 effector complex, which leads to target cleavage and cessation of cOA production, this ensures termination of the antiviral response upon invader clearance. In some type III CRISPR-Cas systems, the ring nuclease activity may limit the self-toxicity of Csm6 enzymes, particularly in systems that do not possess standalone ring nucleases. Consistent with this, we demonstrate that impairment of the CARF domain ring nuclease activity in EiCsm6 results in its hyperactivation and CRISPR self-toxicity. Finally, the widespread occurrence of type III CRISPR-Cas systems, as well as the recently discovered bacterial cGAS/DncV-like nucleotidyltransferase (CD-NT) enzymes that synthesize diverse cyclic di-and trinucleotides^28^, combined with the diversity of their downstream effectors^29,30^, suggests that cyclic oligonucleotide-dependent signaling is an important aspect of prokaryotic genome defense systems and awaits further mechanistic studies of their regulation.

## Methods

### Protein expression and purification

The gene sequence of EiCsm6 was amplified from genomic DNA (*Enterococcus italicus* DSM-15952) by PCR and cloned into a 1B vector (Addgene 29653) using ligation-independent cloning resulting in constructs carrying an N-terminal hexahistidine tag followed by a Tobacco Etch Virus (TEV) protease cleavage site. Point mutations were introduced by inverse PCR and verified by DNA sequencing. EiCsm6 and its mutants were purified similarly as described before^11^. Briefly, EiCsm6 was expressed in BL21 Star (DE3) E. coli cells grown in LB medium by induction with 0.5 mM IPTG at 18 °C for 16 h. Harvested cells were resuspended in lysis buffer (20 mM HEPES pH 7.5, 500 mM KCl and 5 mM imidazole), lysed by sonication, and the lysate was clarified at 20000 x*g* for 30 min. The clarified lysate was loaded onto a 5 ml Ni-NTA HisTrap (GE Life Science) column. The column was washed with lysis buffer supplemented with increasing concentrations of imidazole and the protein was eluted with 20 mM HEPES pH 7.5, 500 mM KCl and 250 mM imidazole. The elution fractions were pooled and diluted with SEC buffer (20 mM HEPES pH 7.5, 500 mM KCl) to a final imidazole concentration of 50 mM. To remove the hexahistidine tag, the sample was incubated at 4 °C for 16 h in the presence of His-tagged TEV protease. The sample was then reapplied to the Ni-NTA affinity column to remove uncleaved protein and the protease. In a final step, the protein was purified by size exclusion chromatography in SEC buffer using a Superdex S200 16/60 column (GE Healthcare). Csm6-containing fractions were identified by SDS-PAGE and concentrated using a 100,000 Da molecular weight cut-off centrifugal filter (Merck Millipore) to a final concentration of approximately 20 mg ml^-1^ and stored at −80 °C until use.

### X-ray crystallography

Crystals of EiCsm6 were initially grown in the presence of cA6 (2-fold molar excess) using the hanging drop vapor diffusion method. 50 μM of EiCsm6 and 100 μM of cA6 were mixed 1:1 (1 μl + 1 μl) with reservoir solution containing 100 mM Bis Tris Propane pH 5.5-6.5, 20-35% PEG3350 and 200 mM sodium citrate. Crystals were cryoprotected with reservoir solution supplemented with 25% glycerol and flash cooled in liquid nitrogen. Data were collected at the beamline PXIII of the Swiss Light Source (Paul Scherrer Institute, Villigen, Switzerland) and processed with XDS31. The crystals belonged to space group *P*2_1_2_1_2_1_ and contained one Csm6 dimer per asymmetric unit. Data analysis using phenix.xtriage^32^ indicated the presence of a large off-origin peak (68.7% of the origin value) in the Patterson function, indicative of translational non-crystallographic symmetry (NCS). Experimental phases were obtained by the single isomorphous replacement-anomalous scattering (SIRAS) method in autoSHARP^33^ using native data and a single-wavelength anomalous dispersion (SAD) dataset obtained from a crystal soaked with 5 mM thimerosal for 5 min and measured at the Hg L-III absorption edge (wavelength of 1.00767 Å). Density modification of experimental phases in AutoSHARP resulted in an interpretable map which was used for initial model building of the EiCsm6 homodimer. Notably, there was no indication in the density maps for the presence of cA6 bound to the CARF domain, suggesting that the ligand underwent degradation in situ. The refinement of the atomic model stalled at a *R*_free_ value of ∼40%, presumably due to high degree of translational pseudosymmetry, and could not be completed.

EiCsm6 crystals bound to cFA6 (synthesized by Biolog Life Science Institute GmbH & Co. KG) were grown in drops containing 120 μM EiCsm6 and 288 μM cFA6 equilibrating against a reservoir solution containing 100 mM HEPES pH 7.5, 1.6 M ammonium sulfate and 25 mM lithium perchlorate. The crystals diffracted to a resolution of 2.4 Å, belonged to space group *P*1, and contained four EiCsm6 homodimers, related by translational non-crystallographic symmetry, in the asymmetric unit. A native dataset was collected at the beamline PXIII of the Swiss Light Source and processed with autoproc^34^. Phases were calculated by molecular replacement in Phaser^35^, as implemented in Phenix, using the atomic model of the EiCsm6 protomer obtained by experimental phasing. The resulting electron density map showed clear features corresponding to cFA6 bound to the dimeric interface of the CARF domains of EiCsm6. The atomic model of the EiCsm6-cFA6 complex was completed by iterative building in Coot^36^ and refined with Phenix.refine^37^ (**Supplementary Table 1**). A 2m*F*_O_-D*F*_C_ composite omit38 map was calculated in Phenix using default parameters.

### Ribonuclease activity assays

The RNase activity of EiCsm6 and its activation was assayed using a fluorogenic substrate similarly as described before^22^. Briefly, cA6, cFA6 (both synthesized by Biolog Life Science Institute GmbH & Co. KG), or supernatants from cA6 degradation reactions were incubated with EiCsm6 or its mutants in 10 mM HEPES pH 7.5 and 50 mM KCl in a total volume of 100 μl. The reaction was started by addition of 2 pmol (0.8 pmol for the spike-in experiment) of RNaseAlert substrate (Integrated DNA Technologies). Fluorescence signal (excitation at 490 nm, emission at 520 nm) was measured over time using a PHERAstar FSX (BMG Labtech) multimodal plate reader. All experiments were done in triplicates and the data points represent the mean +/-standard error of the mean (s.e.m.). Assays comparing EiCsm6 activation by cA6 and cFA6 (**Supplementary Figure 3**) used 100 nM (final concentration) of ligand and 1 nM of WT EiCsm6. Experiments to compare the effect of CARF domain mutations (**Fig. 2d**) used 10 nM of cA6 and 0.5 nM of each EiCsm6 mutant. For the RNase substrate spiking experiment (**Fig. 3b**), WT EiCsm6 or the T11A mutant (10 nM final concentration) was mixed with cA6 (500 nM final concentration) and 0.8 pmol of RNaseAlert substrate at the start of the reaction, and additional RNaseAlert substrate (0.8 pmol) was spiked-in every 5 minutes.

For the pre-incubation assay of cA6 with EiCsm6 or its mutants (**Supplementary Figures 2** and **7**), 100 nM of protein were incubated with 1 μM of cA6 with for 60 min at 37 °C in a total volume of 100 μl. The reaction was stopped by incubating the sample at 95 °C for 5 min and denatured EiCsm6 was removed by centrifugation (16000 x*g* for 5 minutes). Subsequently, 2 μl of the reaction supernatant (or 20 nM of cA6 as positive control) were added to 2 nM of WT EiCsm6 for activity assay using RNaseAlert as described above. The experiments were done in parallel and the results are presented in two figures for clarity, with the same set of controls. For the pre-incubation assay with cA6 or cFA6 (**Supplementary Figure 3**), 1 μM of WT EiCsm6 was incubated with 10 μM of cA6 or cFA6 for 60 min at 37 °C in a total volume of 100 μl. After inactivation, 1 μl of the reaction supernatant (or 0.1 μM cA6 as positive control) were used with 20 nM WT EiCsm6 in the RNaseAlert activity assay.

### Csm6 toxicity growth curves

Growth curves from EiCsm6 interference against pTarget was performed as previously described^17^. Briefly, overnight cultures containing plasmids pTarget and pCRISPR (pJTR366, pJTR367, pJTR368, pJTR369, pJTR417, pJTR422, or pGG-BsaI-R for the dHD loss-of function experiments, and pJTR413, pJTR415, or pGG-BsaI-R for the gain-of-function experiments) were diluted and normalised for optical density (OD) measured at a wavelength of 600 nm. Cells were then seeded in a 96 well plate and target transcription was induced with anhydrous tetracycline (12.5 ng ml^-1^ for the experiment shown in Fig. 2e, or 5 ng ml^-1^ for the experiment shown in Fig. 3c). OD readings were then obtained every 10 minutes with a microplate reader (TECAN Infinite 200 PRO).

### Molecular cloning for in vivo assays

pJTR366 was generated by amplifying pJTR109 with JTR632 and JTR633, and pJTR179 with JTR915 and JTR918), and combining the PCR products by Gibson assembly^39^. pJTR367 was generated by amplifying pJTR109 with JTR632 and JTR633, and pJTR181 with JTR915 and JTR918), and combining the PCR products by Gibson assembly. pJTR368 was generated by first making pJTR355 (by amplifying pJTR179 with JTR223 and JTR924, and with JTR921 and JTR224, and combining the PCR products by Gibson assembly), and them amplifying pJTR109 with JTR632 and JTR633 and pJTR355 with JTR915 and JTR918, and combining the PCR products by Gibson assembly. pJTR369 was generated by first making pJTR356 (by amplifying pJTR179 with JTR223 and JTR924, and by JTR921 and JTR224, and combining the PCR products by Gibson assembly), and amplifying pJTR109 with JTR632 and JTR633, and pJTR356 with JTR915 and JTR918, and combining the PCR products by Gibson assembly. pJTR417 was generated by amplifying pJTR366 with W852 and JTR967, and with W614 and JTR966, and combining the PCR products by Gibson assembly. pJTR422 was generated by amplifying pJTR366 with W852 and JTR969, and with W614 and JTR968, and combining the PCR products by Gibson assembly. pJTR413 was generated by amplifying pJTR360 with W852 and PS466, and with W614 and PS465, and combining the PCR products by Gibson assembly. pJTR415 was generated by amplifying pJTR366 with W852 and PS466, and with W614 and PS465, and combining the PCR products by Gibson assembly. pGG-BsaI-R was described previously^40^. The plasmids and primers used for in vivo assays are described in **Supplementary Tables 2** and **3** respectively.

### EC_50_ analysis for cFA6

RNase activities were measured for reactions containing 5 nM of EiCsm6 supplemented with the indicated cFA6 concentrations. Initial velocities were calculated during the first 100 s and were plotted against cFA6 concentrations in GraphPad and fitted by using a nonlinear regression analysis using the log(dose)-versus-response relationship, assuming a variable Hill coefficient.

### LC-MS analysis

The degradation of cA6 or cFA6 was assayed by incubating equimolar amounts (final concentration of 25 μM) with WT EiCsm6 or the H377A mutant at 37 °C for 1 hour in a total volume of 40 μl for Supplementary Figures 1 and 3. For Supplementary Figure 6a, 5 μM of EiCsm6 T11A/R372A/N373A/H377A or R372A/N373A/H377A were incubated with 50 μM of cA6 at 37 °C for 30 minutes. The reaction was stopped by incubating the sample at 95 °C for 5 min. The denatured protein was removed as described above. LC-MS analysis was performed on an Agilent 1200/6130 LC-MS equipped with a Waters Acquity UPLC OST C18 column (2.1×50 mm, 1.7 μm) at 65 °C. 30 μl of sample were injected. Eluent A was aqueous hexafluoroisopropanol (0.4 M) containing triethylamine (15 mM). Eluent B was methanol. Elution parameters are provided in **Supplementary Table 4**.

### cFA6 synthesis

All reagents and solvents for chemical operations were of analytical grade or the highest-purity grade available from commercial suppliers. Solvents for chromatographic operations were specified as analytical grade, HPLC-grade, or gradient HPLC-grade. YMC*Gel SIL (6 nm, S-75 µm) was used for preparative flash chromatography and TLC was performed with Merck 60 F254 silica gel plates. All chromatographic operations were performed at ambient temperature. Evaporation of solvents was accomplished by rotary evaporation *in vacuo* either with membrane pump vacuum or oil pump high vacuum with water bath temperatures not exceeding 30–33°C.

UV-spectra were recorded on a JASCO V-650 spectrometer in phosphate buffered aqueous solution (pH 7). Mass spectra were generated with a Bruker Esquire LC 6000 spectrometer in the electrospray ionization mass spectrometry (ESI-MS) mode with 50% water/49.5% methanol/0.5% NH_3_ (pH 9-10) as matrix. Nuclear magnetic resonance (NMR) spectra were recorded with a 400 MHz Bruker Avance III HD and chemical shifts are expressed in parts per million (ppm). Chemical shifts were referenced to the DMSO solvent signal, 2.50 ppm for ^1^H. 85% phosphoric acid was used as external standard for ^31^P NMR spectra with 0 ppm. All ^31^P NMR spectra were recorded with proton decoupling.

Analytical HPLC was performed using a VWR / Hitachi: L-7100 Pump; VWR / Hitachi: L-7400 variable wavelength UV/Vis detector and VWR / Hitachi: D-7500 Integrator.

For the chemical synthesis 2.3 mmol of cyanoethyl phosphoramidite 5’-DMTr-2’-F-3’-CEP-N^6^-Bz-adenosine (ChemGenes, Wilmington, MA, USA, Cat. No. ANP-9151) were used as starting material for the stepwise synthesis of the protected linear hexanucleotide precursor 5’-OH-2’-F-N^6^-Bz-adenosine-(3’→5’)-cyanoethyl-phosphate-2’-F-N^6^-Bz-adenosine-(3’→5’)-cyanoethyl-phosphate-2’-F-N^6^-Bz-adenosine-(3’→5’)-cyanoethyl-phosphate-2’-F-N^6^-Bz-adenosine-(3’→5’)-cyanoethyl-phosphate-2’-F-N^6^-Bz-adenosine-(3’→5’)-cyanoethyl-phosphate-2’-F-3’-H-phosphonate-N^6^-Bz-adenosine with a standard oligonucleotide coupling protocol^41^ with the following modifications: After initial phosphonate preparation and after each coupling step with 1.4 eq. 5’-DMTr-2’-F-3’-CEP-N^6^-Bz-adenosine and 2 eq. 5-ethylthio-tetrazole as coupling reagent the raw material was purified by preparative flash chromatography on silica gel using a step-gradient of methanol in dichloromethane. The final cyclization step and the release of protection groups was performed according to a standard protocol^41^, leading to the raw product cFA6 after evaporation of solvents. 100 mL water was added and the resulting suspension was placed in an ultrasonic bath at room temperature for 15 minutes. Afterwards, additional 100 mL water was added followed by 3 extraction cycles with 200 ml chloroform each. The combined organic phases were extracted with 300 ml water and the combined product-containing aqueous phase was filtered with a 0.45 µm regenerated cellulose (RC) filter and concentrated under reduced pressure. The complex reaction mixture was initially purified by repeated preparative reversed phase medium pressure liquid chromatography (MPLC). The product solution was applied to a Merck LiChroprep®RP-18 column (15 - 25 µm; 450 × 50 mm) previously equilibrated with 100 mM triethylammonium formate (TEAF, pH 6.8) in water. Elution was performed with a step-gradient of 3%, 4%, 6%, 8% and 10% 2-propanol, 20 mM TEAF (pH 6.8) in water. Subsequent purifications of product containing fractions were accomplished by a Merck LiChroprep®RP-18 column (15 - 25 µm; 450 × 25 mm) with 2-propanol as organic modifier. Final purifications were performed by semi-preparative reversed phase HPLC with a YMC*GEL ODS-A 12 nm column (10 µm; 250 × 16 mm), previously equilibrated with 100 mM sodium dihydrogen phosphate (NaH_2_PO_4_) buffer (pH 6.8). Elution was performed with a step-gradient of 8%, 9% and 10% acetonitrile, 50 mM NaH_2_PO_4_ (pH 6.8) in water. For desalting, cFA6 fractions of sufficient purity were applied to a Merck LiChroprep® RP-18 column (15-25 µm; 450 × 25 mm), previously equilibrated with water. The column was washed with water to remove excess TEAF or NaH_2_PO_4_ buffer. Afterwards, 2% 2-propanol in water was used to elute the desalted cFA6. To generate the sodium salt form of cFA6, pooled product-containing fractions were partially concentrated under reduced pressure and subsequently applied to a Toyopearl™ SP-650M cation exchange column (65 µm; 100 × 10 mm) Na^+^-form, previously regenerated with 2 M sodium chloride and washed with water. For elution the column was washed with water until no UV-absorbance was detectable at 254 nm anymore. After filtration and careful evaporation under reduced pressure, 36.65 µmol cFA6, sodium salt was isolated with a purity of 99.66% by HPLC (theoretical yield: 1.59%).

Compound name: 2’-,2’’-,2’’’,2’’’’,2’’’’’,2’’’’’’-hexadeoxy-2’-,2’’,2’’’,2’’’’,2’’’’’,2’’’’’’-hexafluoro-cyclic hexaadenosine monophosphate (cFA6; 2’,2’’,2’’’,2’’’’,2’’’’’,2’’’’’’-hexa-F-c-hexa-dAMP), sodium salt

Formula (free acid): C_60_H_66_F_6_N_30_O_30_P_6_ (MW 1987.18 g/mol)

UV-Vis (water pH 7.0): λ_max_ 259 nm; ε 81000.

ESI-MS pos. mode: m/z 1071 (M+H+7 Na)^2+^, m/z 1060 (M+H+6 Na)^2+^,

ESI-MS neg. mode: m/z 992 (M-H)^2-^, m/z 1986 (M-H)^-^.

^1^H NMR (400 MHz, D_2_O) δ 8.12 (s, 6H), 7.91 (s, 6H), 6.18 (dd, *J* = 15.7, 3.2 Hz, 6H), 5.54 (ddd, *J* = 51.1, 4.7, 3.3 Hz, 6H), 5.08-4.95 (m, 6H), 4.56 (s, 6H), 4.41 – 4.24 (m, 12H) ppm.

^31^P NMR (162 MHz, D_2_O): δ −0.31 (s, 6P) ppm.

Analytical HPLC (Kromasil 100-10, RP-8 (10 µm; 250 × 0.4 mm)) 8% acetonitrile, 25 mM

NaH_2_PO_4_ buffer, pH 6.8; 1.5 mL/min; UV 260 nm: t_RET_ 10.59 min.

## Data Availability

The data that support the findings of this study are available as Source Data file. The atomic coordinates of EiCsm6 bound to cFA6 been deposited in the Protein Data Bank (PDB) under accession code 6TUG [http://dx.doi.org/10.2210/pdb6TUG/pdb].

## Acknowledgements

We thank Vincent Olieric, Takashi Tomizaki and Meitian Wang (Swiss Light Source, Paul Scherrer Institute, Villigen, Switzerland) for assistance with crystallographic data collection and analysis. We thank members of the Jinek laboratory for helpful discussions and critical reading of the manuscript. This work was supported by the European Research Council (ERC) Consolidator Grant CRISPR2.0 (Grant no. ERC-CoG-820152, M.J.), the Swiss National Science Foundation Grant 205321_169612 (J.H), and the NCCR RNA and Disease. J.T.R. was supported by a Boehringer Ingelheim Fonds PhD fellowship. L.A.M. is supported by a Burroughs Wellcome Fund PATH Award and an NIH Director’s Pioneer Award (DP1GM128184). L.A.M. is an Investigator of the Howard Hughes Medical Institute. M.J. is International Research Scholar of the Howard Hughes Medical Institute and Vallee Scholar of the Bert L & N Kuggie Vallee Foundation.

## Author Contributions

C.G-D. and M.J. designed experiments. C.G-D. prepared samples, determined crystal structure of EiCsm6 and performed nuclease activity and cA6 degradation assays. O.T. and O.N. performed mutagenesis and prepared samples. J.T.R. performed in vivo experiments under the supervision of L.A.M. F.S. synthesized cFA6. C.B. performed LC-MS analysis, under the supervision of J.H. C.G-D. and M.J. wrote the manuscript with input from the other authors.

