## Supplemental Information for "Activation and self-inactivation mechanisms of the cyclic oligoadenylate-dependent CRISPR ribonuclease Csm6"

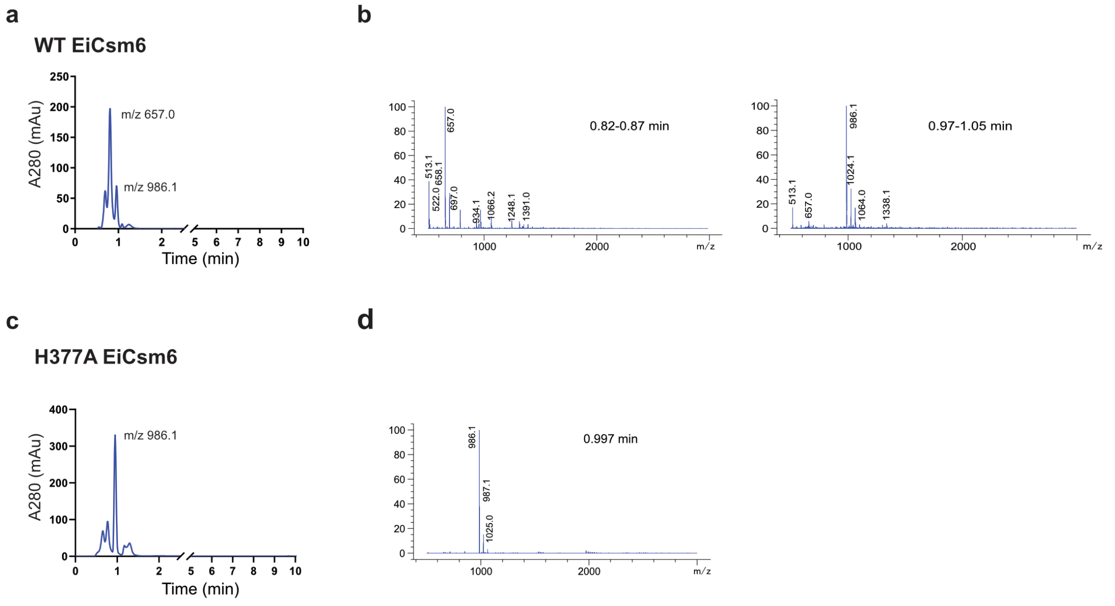

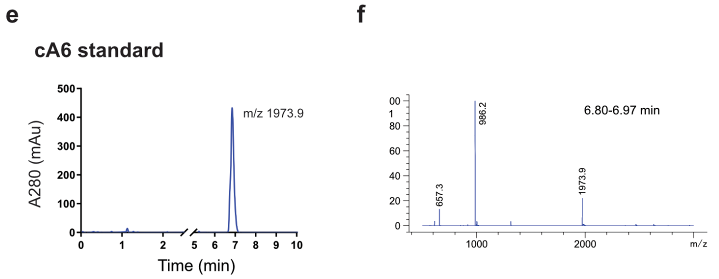

g

| Molecule | Calculated m/z |
|---|---|
| cA6 | 1975.3 |
| cA3/A3>P | 987.6 |
| cA2/A2>P | 658.4 |

**Supplementary Figure 1. Degradation of cA6 by EiCsm6**
(**a, c, e**) LC-MS analysis of the degradation products generated after incubating equimolar amounts (25 μM final concentration) of cA6 with WT EiCsm6. Source data are provided as a Source Data file. (**a**) or H377A HEPN domain EiCsm6 mutant (**c**). (**e**) cA6-only control. (**b, d, f**) Mass spectra of the peaks obtained in **a**, **c**, **e**. Source data are provided as a Source Data file. m/z peak of 986.1 corresponds to a tri-AMP species containing a terminal 2',3'-cyclic phosphate. m/z peak of 657.0 corresponds to a di-AMP species containing a terminal 2',3'-cyclic phosphate. Note that 657.0 m/z peak is only detectable in the WT EiCsm6 sample and absent from the H377A EiCsm6 sample, suggesting that the dinucleotide is a product of HEPN domain-catalyzed degradation. (**g**) Calculated theoretical m/z ratios for the species present in a-f.

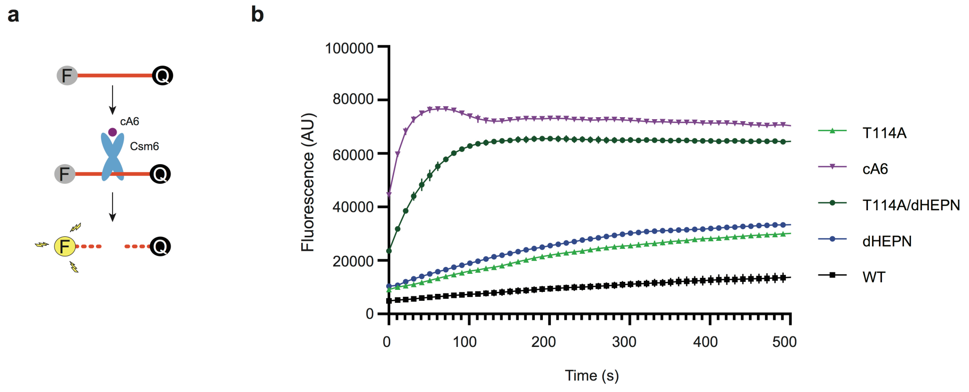

**Supplementary Figure 2. The CARF domain of EiCsm6 is a ring nuclease**
(**a**) Schematic diagram of the fluorogenic ribonuclease activity assay. The RNaseAlert substrate is covalently modified with fluorophore (F) and a quencher (Q) moieties. Substrate cleavage leads to de-quenching of the fluorophore and is detected as an increase in fluorescence. (**b**) Fluorogenic RNase activity assay of WT EiCsm6 (2 nM) activated by degradation products obtained by pre-incubating cA6 (1 μM) with WT, R372A/N373A/H377A (dHEPN) EiCsm6, T114A/R372A/N373A/H377A (dHEPN/T114A) EiCsm6 proteins (100 nM) or without protein (cA6) for 60 min at 37 °C. Data points represent the mean of three replicates; error bars represent the standard error of the mean (s.e.m.). AU, arbitrary units. Source data are provided as a Source Data file.

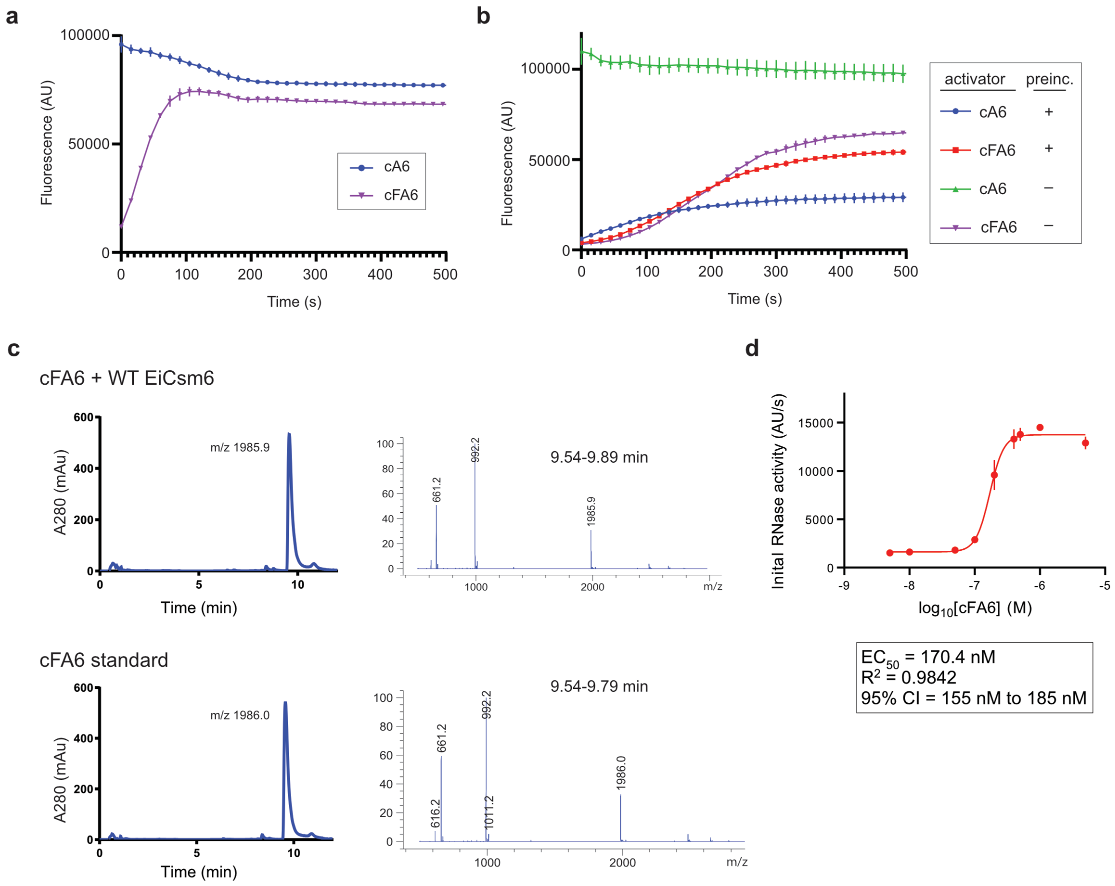

**Supplementary Figure 3. cFA6 activates EiCsm6**

(**a**) RNase activity assay of WT EiCsm6 (1 nM) activated with cA6 or cFA6 (100 nM final concentration). Source data are provided as a Source Data file. (**b**) RNase activity assay of WT EiCsm6 activated with cA6 or cFA6 pre-incubated in the presence or absence of WT EiCsm6. Source data are provided as a Source Data file. (**c**) LC-MS analysis and mass spectra of the products obtained after incubating WT EiCsm6 with cFA6 (top) and cFA6 control (bottom). The calculated m/z for cFA6 is 1987.2. (**d**) EC50 analysis of cFA6 by plotting RNase activity (initial velocity) as a function of cAF6 concentration. AU, arbitrary units. Data points represent the mean of three replicates; error bars represent the standard error of the mean (s.e.m.). Source data are provided as a Source Data file.

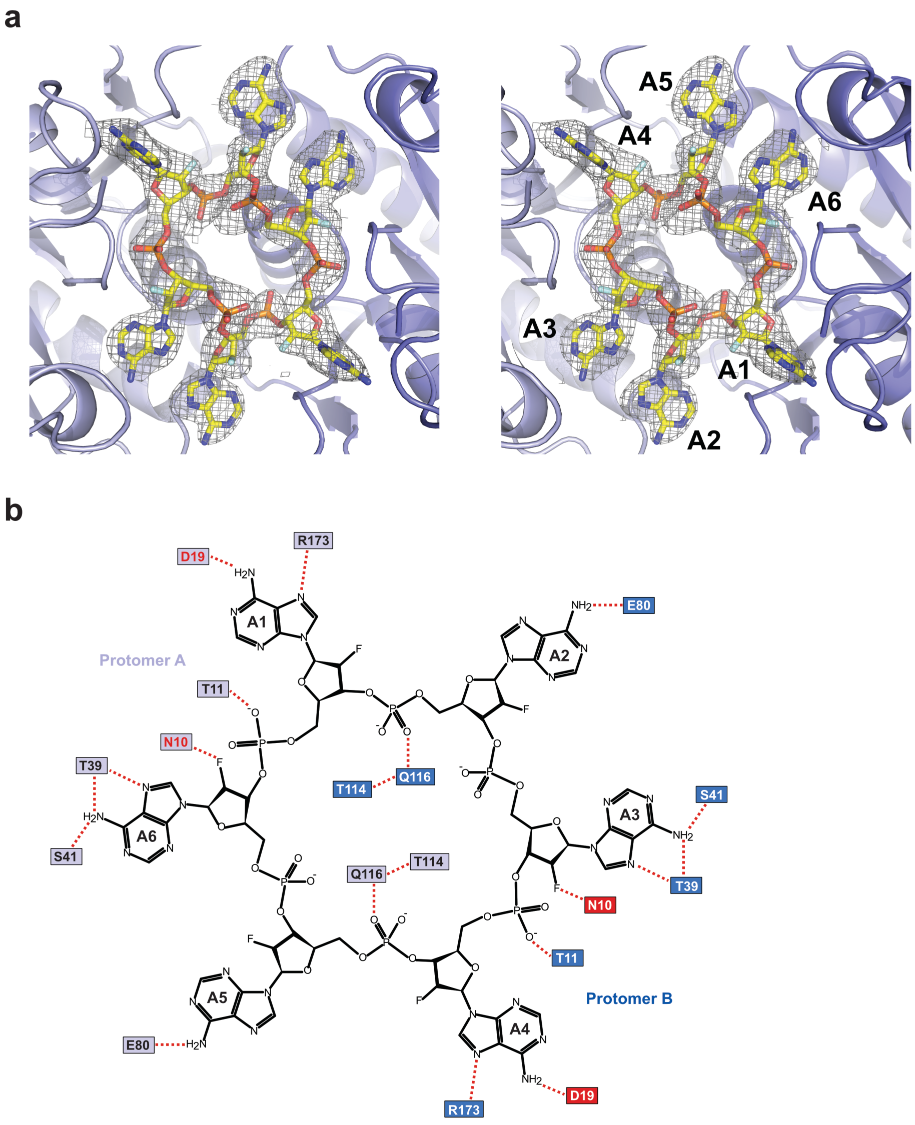

**Supplementary Figure 4. Molecular recognition of cFA6 by EiCsm6**
(**a**) A zoom-in stereo view of the cFA6 ligand bound at the CARF domain interface of EiCsm6. The electron density represents a 2m$F_O$-D$F_C$ composite omit map, contoured at 1.0 σ and displayed within a radius of 2.2 Å around cFA6. (**b**) Schematic representation of EiCsm6-cFA6 interactions. Hydrogen-bonding contacts are depicted with dashed red lines. Amino acid residues contacting cFA6 via the peptide backbone are highlighted in red.

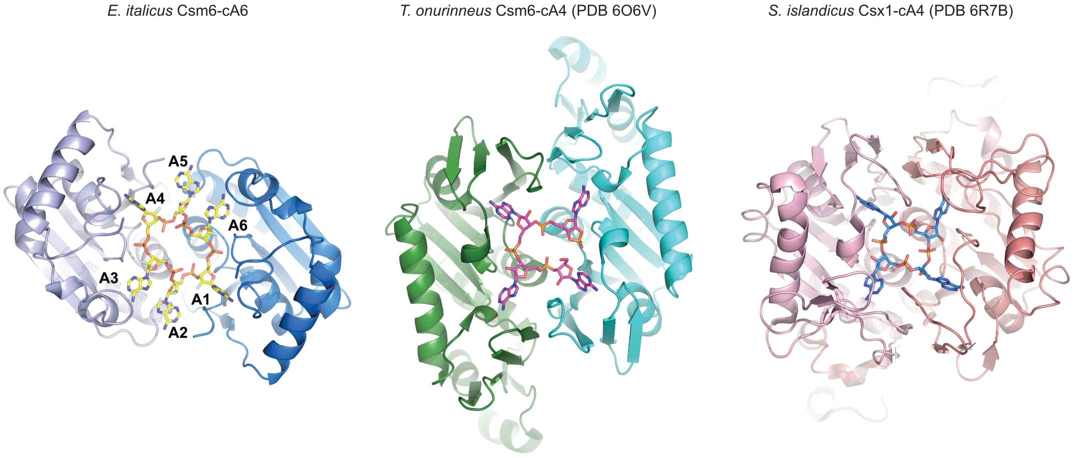

**Supplementary Figure 5. Structural comparison of CARF domains**
Structural comparisons of the cOA binding sites in the EiCsm6-cFA6, *T. onnurineus* Csm6-cA4 and *S. islandicus* Csx-cA4 complexes. The structure were superimposed in Coot[36] and are shown in the same orientation

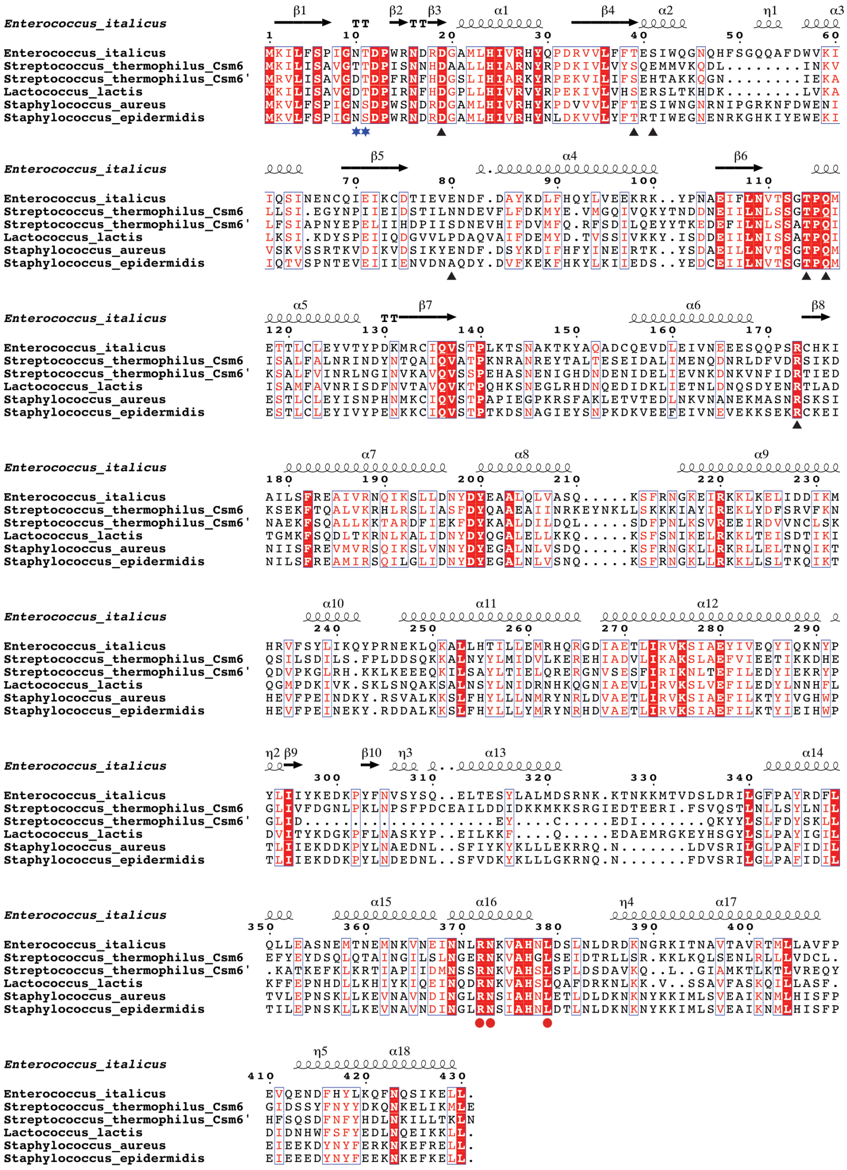

**Supplementary Figure 6. Multiple sequence alignment of Csm6 orthologs**

Multiple sequence alignment of Csm6 orthologs from the type III CRISPR-Cas systems of *Enterococcus italicus* (WP_007208953.1), *Streptococcus thermophilus* (WP_014621552.1 and WP_014621551.1), *Lactococcus lactis* (AGA14268.1), *Staphylococcus aureus* (EZS13226.1) and *Staphylococcus epidermidis* (WP_002502662.1). The alignment was generated using Clustal Omega[42] and displayed using Espript[43]. The secondary structure of EiCsm6 is schematically depicted above the sequences. CARF domain residues involved in cA6 binding are marked with black triangles. Residues implicated in cA6 degradation are marked with blue stars. Catalytic residues in the HEPN RNase domain are marked with red dots.

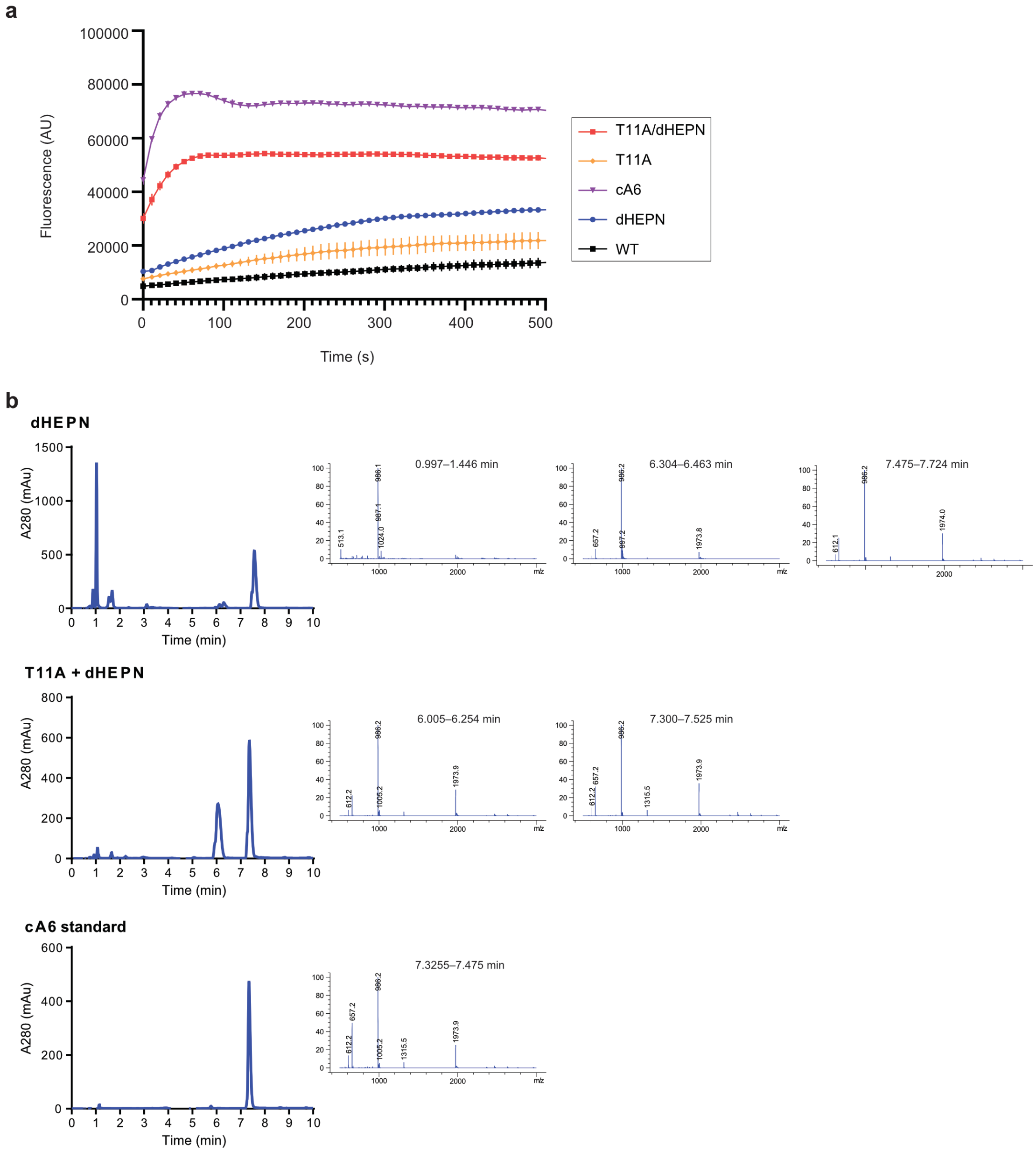

**Supplementary Figure 7. Degradation of cA6 by EiCsm6**
(**a**) Fluorogenic RNase activity assay of WT EiCsm6 (2 nM) activated by degradation products obtained by pre-incubating cA6 (1 μM) with WT, R372A/N373A/H377A (dHEPN) EiCsm6, T11A, T11A/R372A/N373A/H377A (T11A/dHEPN) EiCsm6 proteins (100 nM) or without protein (cA6) for 60 min at 37 °C. Data points represent the mean of three replicates; error bars represent the standard error of the mean (s.e.m.). AU, arbitrary units. The data were generated in the same experiment as data in Supplementary Figure 2 and, are shown separately for clarity, with the same set of controls (cA6, WT, dHEPN). Source data are provided as a Source Data file. (**b**) LC-MS analysis of the degradation products generated after incubating equimolar amounts (25 μM final concentration) of cA6 with WT EiCsm6 or T11A EiCsm6. Source data are provided as a Source Data file.

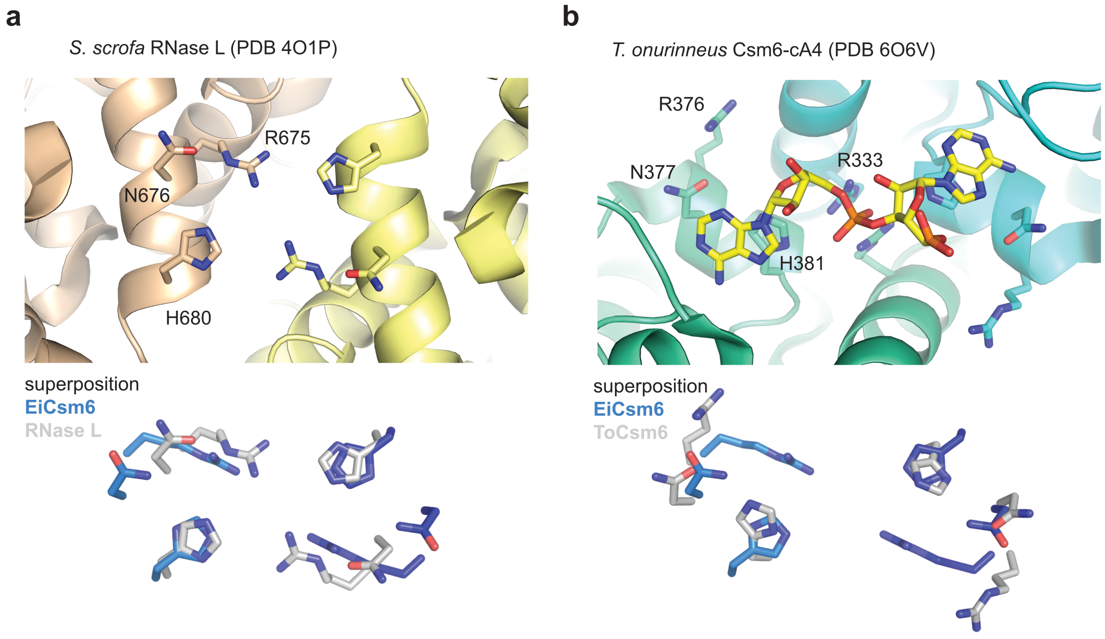

**Supplementary Figure 8. HEPN domain ribonuclease active site of EiCsm6**
Structures of the active forms of porcine RNase L (PDB 4O1P, left) and *Thermococcus onnurineus* Csm6 (PDB 6O6V, right). Bottom panels show superpositions of the conserved HEPN active site residues of RNase L (left) and ToCsm6 (right) with the EiCsm6 structure.

**Supplementary Table 1. Crystallographic data collection and refinement statistics**

| Dataset | EiCsm6–cFA6 | EiCsm Hg SIRAS |
|---|---|---|
| X-ray source | SLS PXIII | SLS PXIII |
| Space group | $P1$ | $P2_12_12_1$ |
| Cell dimensions | | |
| *a, b, c* (Å) | 93.60, 106.69, 116.79 | 58.68, 103.49, 219.71 |
| *α, β, γ* (°) | 90.03, 90.01, 90.10 | 90.00, 90.00, 90.00 |
| Wavelength (Å) | 1.00000 | 1.00767 |
| Resolution (Å)* | 48.51-2.42 (2.51-2.42) | 43.81-2.8 (2.9-2.8) |
| $R_{sym}$ (%)* | 7.65 (43.34) | 23.5 (142.0) |
| *CC1/2 (%)* | 99.8 (78.1) | 100.0 (99.1) |
| $I/\sigma I$* | 12.14 (2.47) | 27.87 (5.83) |
| Completeness (%)* | 90.8 (97.8) | 98.72 (99.137) |
| Redundancy* | 3.5 (3.7) | 106.5 (105.3) |
| **Refinement** | | |
| Resolution (Å) | 48.51-2.42 | |
| No. reflections | 156268 | |
| $R_{work}/R_{free}$ | 0.2635/0.2917 | |
| **No. atoms** | | |
| Protein | 27730 | |
| cFA6 ligand | 528 | |
| Water | 345 | |
| **B-factors** | | |
| mean | 50.78 | |
| Protein | 51.15 | |
| cFA6 ligand | 38.76 | |
| Water | 38.29 | |
| **R.m.s. deviations** | | |
| Bond lengths (Å) | 0.004 | |
| Bond angles (°) | 0.79 | |
| **Ramachandran plot** | | |
| % favoured | 98.67 | |
| % allowed | 1.33 | |
| % outliers | 0.0 | |
| **Molprobity** | | |
| Clashscore | 7.55 | |

* Values in parentheses correspond to the highest-resolution shell

### Supplementary Table 2. Plasmid list for in vivo assays

| | |
|---|---|
| pGG-BsaI-R | No spacer control |
| pJTR366 | dHD, EiCsm6 WT |
| pJTR367 | dHD, EiCsm6 Q116A |
| pJTR368 | dHD, EiCsm6 N10A |
| pJTR369 | dHD, EiCsm6 T11A |
| pJTR413 | dCsm3 |
| pJTR415 | dCsm3, T11A |
| pJTR417 | dHD, EiCsm6 S41A |
| pJTR422 | dHD, EiCsm6 T114A |

### Supplementary Table 3. Primer list for in vivo assays

| | |
|---|---|
| JTR223 | CAGCGCATCACACGCAAAAAGG |
| JTR224 | CATCTCAAATTTCGCATTTATTCCAATTTCC |
| JTR632 | ATGATAAATAAAATTACAGTAGAGTTAGACTTGC |
| JTR633 | TATAGCACCTCATTATTTAACTCTTGAAAAC |
| JTR915 | CAAGAGTTAAATAATGAGGTGCTATAATGAAAATACTCTTTAGTCCAATTGG |
| JTR918 | CTAACTCTACTGTAATTTTATTTATCATAGTAGCTCCTTAATTGATTGATTAAATTG |
| JTR921 | CTTTAGTCCAATTGGAgcTACCGATCCTTGGAGAAACG |
| JTR924 | CCAAGGATCtGcATTTCCAATTGGACTAAAGAGTATTTTC |
| JTR966 | TATCTGGCAAGGGAATCAACATTTTTC |
| JTR967 | GAAAAATGTTGATTCCCTTGCCAGATAgcTTCGGTAAAAAACAGTACCACGCG |
| JTR968 | CACCACAGATGGAGACAACTTTGTG |
| JTR969 | CACAAAGTTGTCTCCATCTGTGGTGcACCACTAGTCACATTTAGAAAAATTTCTGC |
| PS465 | GAATCTAGTATGATTGGAGCAATTGCTTCTCCTGTAGTTAGAGATTTGCAAACC |
| PS466 | GGTTTGCAAATCTCTAACTACAGGAGAAGCAATTGCTCCAATCATACTAGATTC |
| W614 | GGTTATACTAAAAGTCGTTTGTTGG |
| W852 | CCAACAAACGACTTTTAGTATAACC |

### Supplementary Table 4. LC-MS analysis

| Time (min) | Eluent A (%) | Eluent B (%) | flow rate (ml $min^{-1}$) |
|---|---|---|---|
| 0 | 99 | 1 | 0.3 |
| 2.0 | 99 | 1 | 0.3 |
| 12.0 | 85 | 15 | 0.3 |
| 12.5 | 5 | 95 | 0.3 |
| 13.0 | 5 | 95 | 0.3 |
